# Hierarchical Modelling of Crossing Fibres in the White Matter

**DOI:** 10.1101/2023.05.24.542138

**Authors:** Hossein Rafipoor, Frederik J. Lange, Christoph Arthofer, Michiel Cottaar, Saad Jbabdi

## Abstract

While diffusion MRI is typically used to estimate microstructural properties of tissue in volumetric elements (voxels), more specificity can be obtained by separately modelling the properties of individual fibre populations within a voxel. In the context of cross-subjects modelling, these so-called fixel-based analyses require identifying equivalent fibre populations. This is usually done post-hoc, after estimating fibre orientations for individual subjects independently and subsequently matching the fixels between subjects. This approach can fail due to individual differences in fibre orientation distributions.

Here, we introduce a hierarchical framework for fitting crossing fibre models to diffusion MRI data in a population of subjects. This hierarchical setup guarantees that the crossing fibres are consistent by construction and, therefore, comparable across subjects. We propose an expectation-maximisation approach to fit the model, which can scale to large numbers of subjects. This approach produces a crossing-fibre white matter fibre template, which can be used to estimate fibre-specific parameters that are consistent across subjects and, hence, can be used in fixel-based statistical analyses.

## INTRODUCTION

Diffusion MRI is widely used to study between-subjects variations in white matter, both in healthy cohorts and in patient populations(Johansen-Berg and T. E. J. Behrens 2014; Jones 2010). Like in voxel-based morphometry for studying grey matter(Ashburner and Friston 2000), comparing white matter between subjects requires care in the between-subjects spatial alignment. This is particularly crucial when the objective of the analysis is the comparison of microstructural effects (from diffusion MRI models), rather than macroscopic variations in gross anatomy.(S. M. Smith et al. 2006)

Recently, fixel-based analysis of white matter has emerged as a technique for cross-subject analysis of white matter not only within each voxel, but also within each fibre population in the voxel(Dhollander et al. 2021; D. A. Raffelt, J.-Donald Tournier, et al. 2017a) In fixel-based analysis, a white matter orientation model is used to assign properties to fibre populations crossing within each voxel (e.g., the density of fibres aligned with a given direction). These metrics are then compared across subjects, allowing investigation of fibre-specific changes. This approach is in contrast to more traditional analyses that use voxelwise metrics, such as fractional anisotropy (FA) or mean diffusivity (MD), as markers of white matter. While these voxelwise metrics have been shown to be highly sensitive to changes in white matter tissue (e.g.(Scholz et al. 2009)), they cannot easily be interpreted in terms of specific changes to the underlying fibre populations within the voxel (see Figure 1).

**Figure 1.**
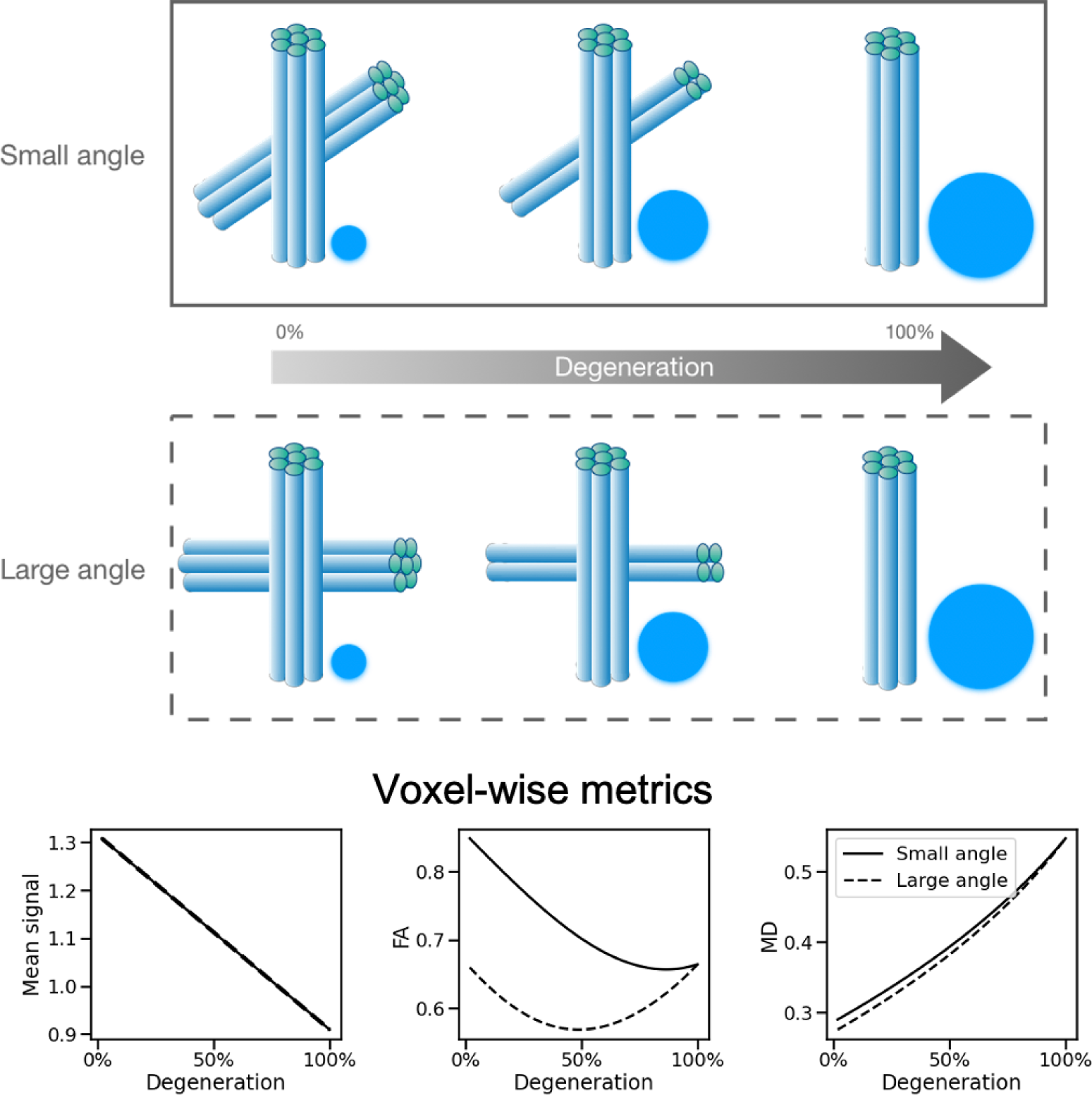
Illustration of the effect of a fibre-specific biological process on voxel-wise measurements. The top panels show a scenario of axonal degeneration in a crossing fibre region, where one of the fibre populations (shown by cylinders) gradually degenerates and is replaced by water with isotropic diffusion (shown by circles). Two possible scenarios for the same process are depicted in the two panels, one in which the angle between the two fibre populations is small, and another in which the angle is large. The bottom row shows the change in voxel-wise measurements of mean signal, fractional anisotropy, and mean diffusivity in both scenarios (solid line for the small angle and dashed line for the large angle). The angle between the fibres has no effect on the mean signal and a small effect on MD, but it substantially affects the changes seen in FA.

Fixel-based approaches attempt to resolve this fibre-specificity issue. By assigning measures to a discrete set of fibre orientations within a voxel, between-subjects differences can then be associated with specific fibre orientations, which in turn may correspond to fibre pathways associated with specific functions. However, not only do fixel-based analyses need to resolve between-subject spatial alignment (just like voxelwise methods do), they also need to resolve between-subject assignment of fibres. In order to “compare like with like”, the discrete set of fibres within a voxel must all be in correspondence across subjects (Jbabdi, T. E. Behrens, and S. M. Smith 2010).

So far, this between-subject fibre assignment has been conducted as a post-processing step, following independent processing of each subject separately. In (Jbabdi, T. E. Behrens, and S. M. Smith 2010), fibre orientations are estimated on a subject by subject basis using the ball and sticks model (T. Behrens et al. 2007). Then, to maximise correspondence both spatially and between subjects, fibre labels are swapped in multiple stages. The method was restricted to the white matter skeleton (S. M. Smith et al. 2006) in order to ensure validity of the spatial assignment. More recently, fixel-based analyses were conducted using continuous fibre ODF models (D. A. Raffelt, J.-Donald Tournier, et al. 2017a; J-Donald Tournier et al. 2019) fitted to each subject separately. Peaks from the fODF were subsequently matched and relabelled across subjects. This approach is, in fact, what is currently referred to as fixel-based analysis.

However, even after accurate spatial normalisation with fibre reorientation, post-hoc between-subjects fibre reassignment can often fail due to variability in white matter orientation estimates. This can either be natural variability in anatomy between subjects, or variability due to noise in the parameter estimation process. In either case, fibre reassignment can be ill-posed.

In this paper, we propose a solution to this problem that takes part earlier in the processing pipeline, at the stage of voxelwise white matter orientation modelling, rendering post-hoc fibre reassignment unnecessary. We build a cross-subject hierarchical model of crossing fibres where, in each voxel, the parameters of the model (including fibre orientations) are drawn from a population distribution. This means that the assignment of the fibre orientations is matched across subjects by construction, rather than through a post-processing step. We fit both the individual subjects and the group parameters simultaneously to all the data and obtain group and individual estimates of orientations and associated fixel parameters. After a template has been created, it can be used to compare signal fractions in different fibres across a larger sample of individuals by fitting the ball-and-sticks model with the template acting as the prior distribution.

In addition, we propose an expectation-maximisation algorithm that enables us to fit the model in a way that is scalable to a large cohort of subjects. We also extend the voxel-based Threshold-Free Cluster Enhancement (TFCE) approach (Smith and Nichols 2009) to enable statistical analysis of fixels with proper correction for multiple comparisons across space. The method and implementation are validated through simulations. Finally, we create a group-averaged fixel template for white matter using data from the UK Biobank (Miller et al. 2016), and we used this template in an example analysis of fixel changes with ageing.

## THEORY

## Crossing fibre model

While our framework can be adapted to work on different crossing fibre models, we choose the ball-and-sticks model because it is simple, widely used, and it lends itself naturally to a hierarchical extension. According to this model, the diffusion signal in a voxel originates from the combination of a ball compartment that represents water with isotropic diffusion and one or more stick compartments that represent water that diffuses along a fibre population. The diffusion signal according to the ball and sticks model is generated by

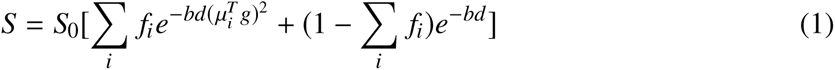

where b, g are the b-value and the gradient orientation, d is the diffusion coefficient, µ*_i_*, *f_i_* are the orientation and signal fraction of the i’th stick and *S*_0_ is the baseline signal with no diffusion encoding. Hence, the set of free parameters in this model are {*S*_0_, *d*, *f_i_*, µ*_i_*} per subject per voxel.

### Hierarchical model

In a hierarchical framework, we assume that, in each voxel, each subject’s parameter (ν*_s_*) represents a sample drawn from a population distribution. This hierarchical structure ensures that all parameters, including the labelling of the fibres, are consistent across subjects.

The population distributions of all scalar parameters are assumed to be Gaussian with unknown means and variances, while orientation parameters are assumed to follow a Watson distribution with unknown mean and concentration. Finally, each subject’s diffusion MRI signal is derived from the subject parameters following the forward model (*M*) in 1 with additive noise (*D_s_* = *M*(ν*_s_*) + E). The noise distribution is assumed to be a zero-centred Gaussian with a variance (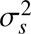). This variance also has an exponential prior distribution with hyperparameter α*_n_*. The diagram in Figure 2 shows the full generative forward model. Using the dependency structure that is shown in the graph, the joint distribution of all free parameters factors into:

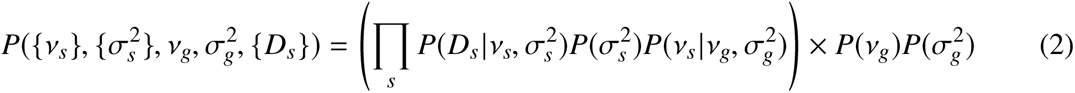

**Figure 2.**
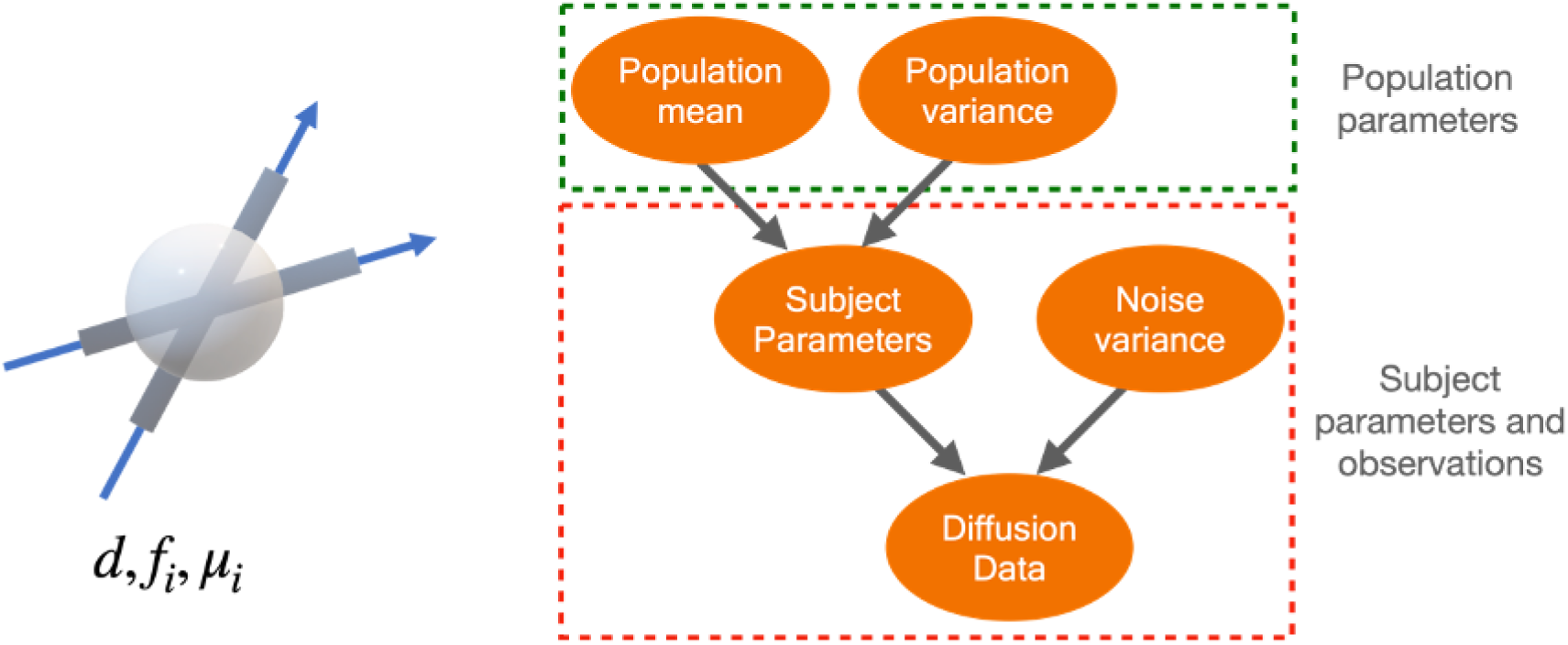
The generative forward model of diffusion MRI signals for the whole population. Left: A schematic representation of the ball and sticks model for diffusion MRI signal in white matter voxels. According to this model, water only diffuses axially inside each population of fibres and isotropically outside the fibres. Right: The hierarchical graphical model for the diffusion MRI signal in a white matter voxel for a population of subjects. In this model, we assume that, within a single subject, each parameter of the ball and stick model is drawn from a population distribution. The red box shows subject-specific parameters and observations, and the green box shows population-level parameters. This hierarchical structure will introduce a dependency between the subjects, which has the advantage of ensuring that the fibre labels are consistent across subjects, therefore allowing for the comparison of fibre-specific parameters between subjects.

The curly brackets indicate sets of subject-specific parameters. The conditional probabilities in the graphical model are defined as:

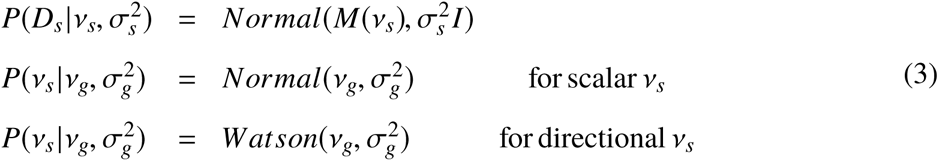

For the Watson distribution, ν is the mean orientation, and 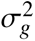 is the orientation dispersion index (H. Zhang, Schneider, et al. 2012). We assume uniform priors within plausible ranges for all the group mean parameters (ν_g_) and a power-law prior for the group variances (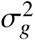) with hyperparameter α_g_. The power law prior is used to prevent the group variance from going to zero. The prior distributions of the parameters are given by

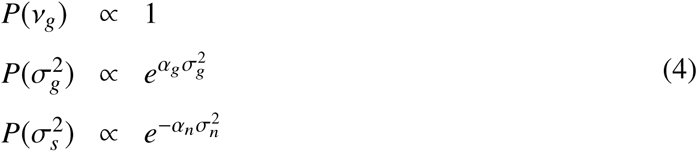

α_g_ and α*_n_*(positive real numbers) are hyperparameters that are adjusted by the user.

### Parameter Inference

We employ a maximum a-posteriori approach to fit the parameters of the hierarchical model to the data, that is to maximise the conditional probability of all the parameters of interest given the data 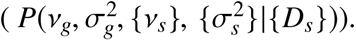 The free parameters of this model are the group means and variances for each model parameter, subject parameters and subject specific noise variance. The full posterior probability of the model according to the hierarchical structure factorises into:

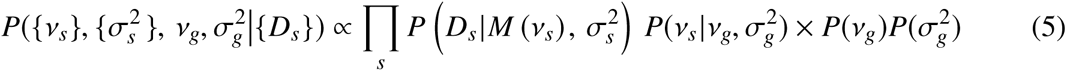

Fitting this entire model requires simultaneously optimising all of these parameters, which creates a space with a very high number of dimensions that grows linearly with the number of subjects. This makes optimization infeasible for datasets with more than a few tens of subjects.

To make the optimisation feasible for large datasets, we employ an expectation-maximisation approach: In the expectation step, the group parameters are estimated while the subject parameters are fixed; in the maximisation step, the group parameters are held fixed while the subject parameters are estimated. These stages are repeated until all of the parameters have converged. Convergence is achieved when all group and subject parameters remain unchanged between subsequent iterations.

### Expectation

In the expectation step, the subject parameters are set to the best guess made so far so that the population parameters can be estimated. Following Equation 2 this leads to optimising the following conditional posterior:

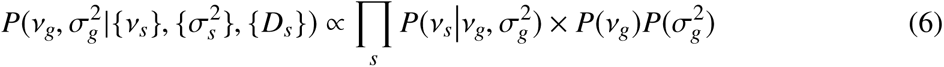

Since we assume that there is no correlation between parameters a-priori, each parameter is evaluated on its own at this stage. For the scalar parameters, plugging the probabilities according to 6 results in:

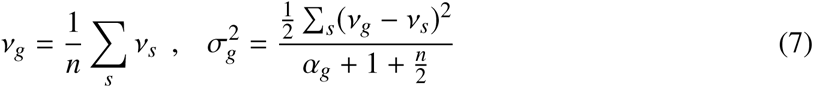

Numerical optimization is used for the maximisation of the posterior to estimate group orientation parameters, including average orientation and dispersion for each fibre population. For all the model fittings (both for individual fits and in the hierarchical model) we have used non-linear model fitting as implemented in the SciPy package Nelder-Mead method. (Nelder and Mead 1965; Virtanen et al. 2020)

### Maximisation

In this stage, the population mean and variance parameters are fixed, and the subject parameters are estimated by maximising the conditional posterior probability. According to Equation 5:

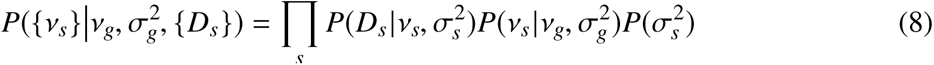

Since the group parameters are fixed in this stage, the likelihood is independent for each subject and factorises into subject specific parts. This makes it possible to optimise each subject’s parameters separately. We therefore independently estimate subject parameters ν*_s_*by optimising the logarithm of the posterior function through numerical optimisation.

### Determining the number of fibres in each voxel

To determine the number of fibres in each voxel, we first fit the hierarchical model using one, two, and three fibres. We then set a threshold on the difference in the quality of the fit achieved by adding a fibre population. The log-likelihood of the data log 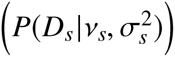 summed over all subjects is used as the quality of fit metric.

Figure 3 shows the full pipeline for fitting the hierarchical model, including the creation of a white matter fixel template representing the group average orientations and population volume fractions in the model.

**Figure 3.**
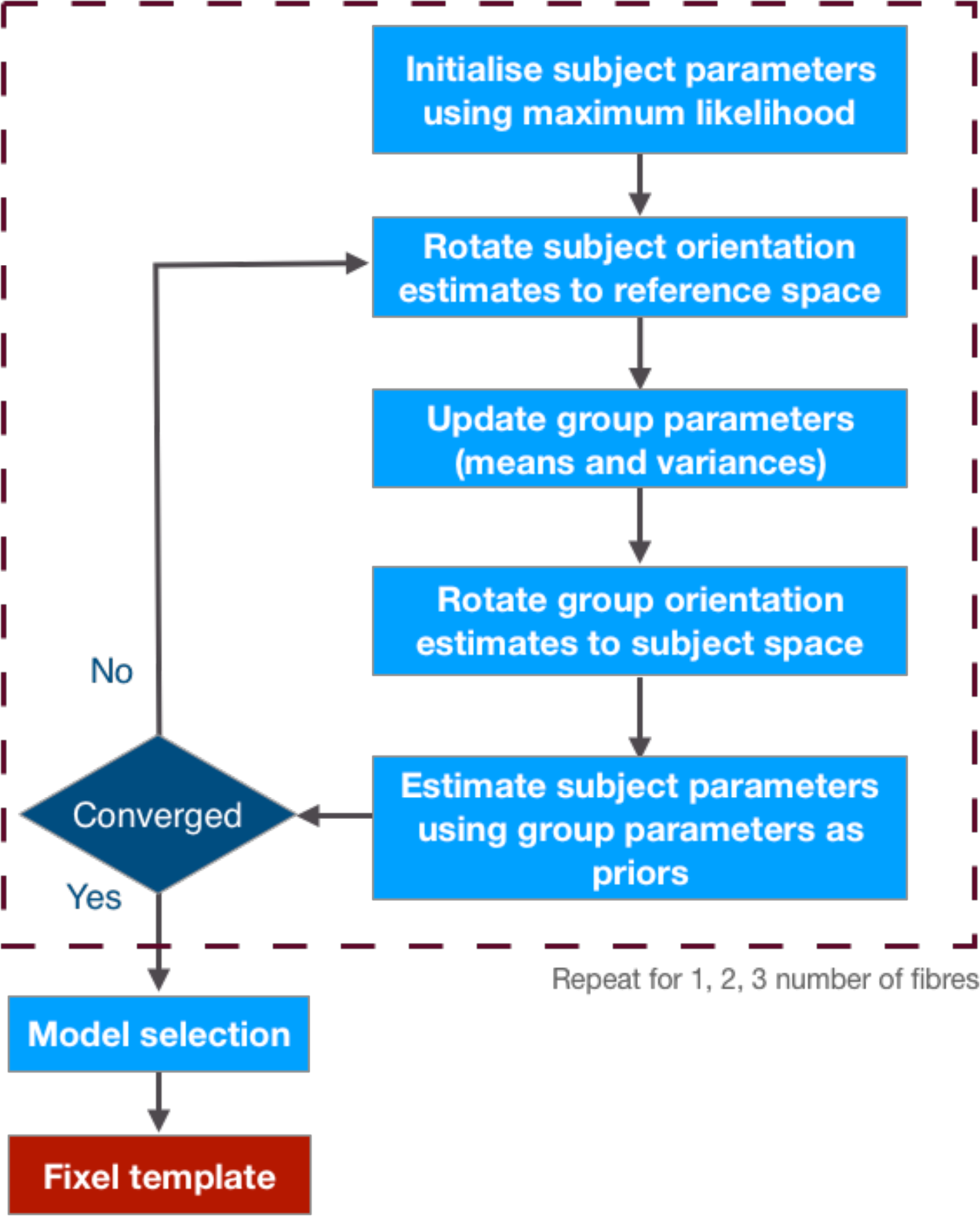
Template creation pipeline. Firstly, the diffusion data is fitted with models for each subject independently using maximum likelihood estimation. We then use the parameters from all subjects to estimate the population distribution for each parameter. We then rotate the orientations back into the space of each subject. The population parameters (mean and variance) are then utilised as prior distributions to re-fit the subject parameter to the diffusion data. This process is repeated until convergence, and for models with 1, 2, and 3 fibres separately. In order to determine the number of fixels within a voxel, we first estimate the improvement in the likelihood function from one to two fibres, and from two to three fibres. Then, we apply thresholds to these improvements to identify the number of fixels in each voxel. This gives us a fixel template that we can use as a prior distribution to fit fixels to each subject. The inputs of the pipeline are diffusion data for each subject that is resampled into a structural template space, and transformations from each subject’s diffusion data to that template. Note that all the computations happen for each voxel in the reference space and all the diffusion data is only fetched once needed, so there is no need to register the diffusion data into the template space. To accomplish this, for every voxel in the reference space we assign a matching voxel from each subject; that is the closest voxel in the subject’s space after transforming the coordinates.

### Fixel based threshold free cluster enhancement

The natural next step following fitting fixels is to fit a cross-subject model of the parameters, e.g., the fibre signal fractions, using a general linear model (GLM). As this is done in each voxel, statistical techniques for controlling false positives are required due to the large number of tests involved. One popular technique is Threshold-Free Cluster Enhancement (TFCE) (Smith and Nichols 2009), which is a method for multiple comparison correction developed for voxelwise cross-subject analyses. TFCE works in an analogous way to cluster-based thresholding methods, but without requiring a cluster-forming threshold. Here, we show how we can extend TFCE to the context of fixel-based analyses.

TFCE works by forming contiguous clusters of voxels that are above a given statistical threshold, then calculating a statistic that combines the cluster height and extent, and finally integrating this statistic across different thresholds. To apply this method to fixel data, we need to modify the definition of spatial neighbourhood used for forming clusters. An example of such modification is found in (D. A. Raffelt, R. E. Smith, et al. 2015), where they utilize tractography to establish the neighborhood structure. In contrast, our approach employs a different rationale for defining the fixel neighborhood.

In our definition, two fixels are regarded as neighbours if they are located in adjacent voxels (27-neighbourhood) and if the angle between them is below a predefined threshold (see Figure 4). This notion of adjacency has three advantages: it is symmetric; it maintains adjacency between fixels that are potentially in the same tract; and, finally, it prevents information from being mixed between crossing fibres within the same voxel. We construct a graph of connections between fixels, where each node in the graph is a fixel, and two fixels that meet the above criteria for being neighbours share an edge.

**Figure 4.**
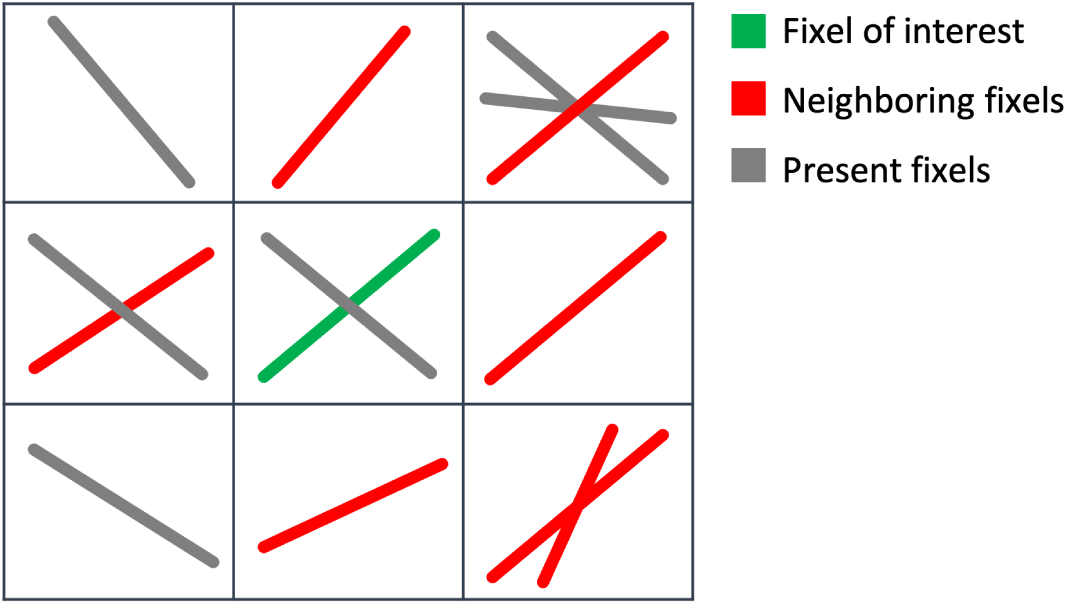
2D section of fixel neighbourhood for TFCE. The neighbours of a fixel (green fixel) are all the fixels that are located in an adjacent voxel (at least one corner is shared, in total 27 voxels) and make angles smaller than a threshold (red fixels). Multiple fixels in a neighbouring voxel may be taken into account as neighbours of the same fixel.

For the fixel of interest, the cluster extent *e*(*h*) at threshold h is defined as the number of nodes in the connected component containing the fixel, only considering the sub-graph that includes fixels with statistics greater than h. As a result, we assign a TFCE score to each fixel as:

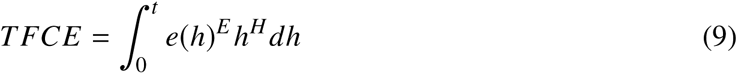

Here E and H are parameters that weigh the importance of cluster extent and height, respectively, *t* is the statistic for the fixel of interest.

Since the TFCE scores may not have a Gaussian distribution, we use standard nonparametric permutation testing to estimate p-values for each fixel. In order to achieve this, we permute the original data labelling (for example, subjects between two groups for examining group differences) and re-estimate the TFCE score multiple times to produce a null distribution. The p-value is then calculated as the likelihood of observing the actual TFCE score or higher in that null distribution.

## METHODS

### Simulation settings

To determine the effectiveness of the hierarchical framework, we evaluated it with simulated data. Diffusion data for 30 participants in a single crossing fibre voxel were simulated using a ball-and-sticks model with two fibres. Each subject’s parameters are randomly sampled from a population distribution. The signal fractions are sampled from Gaussian distributions with means 0.5, 0.3 and standard deviation of 0.1, 0.1 and then truncated to range [0, 1] and normalised to sum to 0.8. Therefore, on average the first fibre population is stronger, but that may not be the case for every single subject. The stick orientations are samples from a Watson distribution with low dispersion (ODI=0.1), so that each fibre population forms a cluster across subjects. Following that, using an acquisition protocol that includes 5 b0 images and 50 diffusion-encoded images with *b* = 1, 2 ms/µm^2^ (identical to that used by the UK biobank dataset (Miller et al. 2016), diffusion data for each subject is simulated using a custom python script. All diffusion signals are then distorted by adding Gaussian noise with SNR=50.

### Data sampling and voxel correspondence

For the hierarchical framework to work, there needs to be spatial correspondence between subjects. This is achieved through image registration of each subjects data to a common reference space. Registration was performed using FSL MMORF (Lange et al. 2020), a multimodal registration tool that uses both scalar structural (T1-weighted) images and diffusion tensor images (DTIs) to drive the registration. We used the Oxford-MultiModal-1 (OMM1) template (Arthofer et al. 2021) as our standard reference space for voxel correspondence. OMM1 is preferred over MNI152 since it was constructed using simultaneous alignment of T1-weighted, T2-FLAIR and DTI images, and therefore includes high quality, unbiased, scalar and tensor data in a common space. The inclusion of DTI data during registration ensures that matching of white matter orientation is taken into account during warp optimization. It’s important to note that using MMORF is not the only option for registration. Alternative registration approaches that incorporate orientations in the alignment process, such as those proposed by (H. Zhang, Yushkevich, et al. 2006), (P. Zhang et al. 2014), are also viable alternatives. This is important for obtaining both correct fixel assignment across subjects, and correct estimates of the group mean and variance of fixel orientation in template space.

We chose not to resample the diffusion data into a shared reference space after registration due to memory and space constraints. Instead, we built correspondence by extracting the necessary data at each stage of inference using the warps between individuals and a reference space, with nearest neighbour interpolation. To move across individual and common spaces, we are required to map the voxel position as well as the registration-induced local re-orientation and apply them to the orientation parameters. Because registration was driven by directional information within the white matter, this step also contributes to reducing cross-subject variability in fibre orientation estimates. In order to rotate all orientation parameters, we used the warpfield’s local Jacobian matrix. We chose not to isolate the rotation component from the Jacobian in order to also account for non-rotational types of distortion, such as shearing. Instead, we used the complete Jacobian matrix to transform the orientation vector and then renormalized the resized orientations to unit length. For example, any given direction parameter in subject native space is resampled to the reference space using:

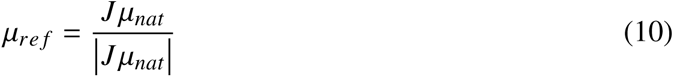

Where µ*_nat_*is the orientation parameter in subject native space, J is the Jacobian matrix of the warp field at that voxel, and µ*_re f_* is the orientation after resampling to the reference space. A very similar approach have been employed in other studies, as demonstrated in (Alexander et al. 2001; Hong, Arlinghaus, and Anderson 2009; D. Raffelt et al. 2012; D. A. Raffelt, J.-Donald Tournier, et al. 2017b).

### Fixel template creation

To create a fixel template, we used the diffusion data of 100 unrelated subjects from the HCP dataset for young adults (Van Essen et al. 2013). The diffusion data includes three shells (b-values: 1, 2, 3 ms/µm^2^ with 90 orientations each) and 18 b0 images (in total 288 volumes). This data has been preprocessed using the standard preprocessing pipeline in FSL for the HCP dataset (Sotiropoulos et al. 2013).

Each subject was registered to OMM1 space using MMORF. Registration was driven using brain-extracted T1-weighted and DTI modalities. As fixel analysis is only valid within white matter, we created a white matter mask by running FSL FAST (Jenkinson et al. 2012; Y. Zhang, Brady, and S. Smith 2001) on the T1-weighted image of the OMM1 template. The template creation process then proceeded as described above. Following the creation of this template, its parameters were fixed and it was used as a prior distribution in order to fit the diffusion model to new, unseen subjects as detailed in the following section.

### GLM analysis

In order to illustrate an example fixel-based analysis utilising the fixel template, we ran a GLM to model changes in fibre signal fractions with age. The diffusion data from 400 randomly selected subjects (200 male, age 50+/-10) in the UK biobank were used for the GLM analysis. This dataset contains two shells (b-values= 1, 2 ms/µm^2^ each in 50 directions) and five b0 images (105 volumes in total). The standard preprocessing pipeline for UK biobank data was used to preprocess the diffusion data (Alfaro-Almagro et al. 2018). Once again, MMORF registration was used to estimate warps for transforming parameters between individuals diffusion space and the OMM1 template space. In this instance T2-FLAIR data were used in addition to T1-weighted and DTI data to drive the registration.

We used fixel signal fractions as dependent variables, and age as the independent variable in our GLM analysis, with sex and head size as confounding factors. The t-statistics for the regression against age were estimated using a linear fit for each fixel independently. Then, we estimated voxel-wise p-values from permutation tests (5000 permutations) using the fixel-based TFCE scores calculated with the proposed cluster enhancement method.

## RESULTS

### Simulations

The individual (independent) fitting approach (maximum likelihood estimates) and hierarchical model are employed to estimate individual subject parameters and population-level distribution parameters from simulated diffusion data with known ground truth. The estimated parameters from each approach versus the actual parameters for each subject is shown in Figure 5.

**Figure 5.**
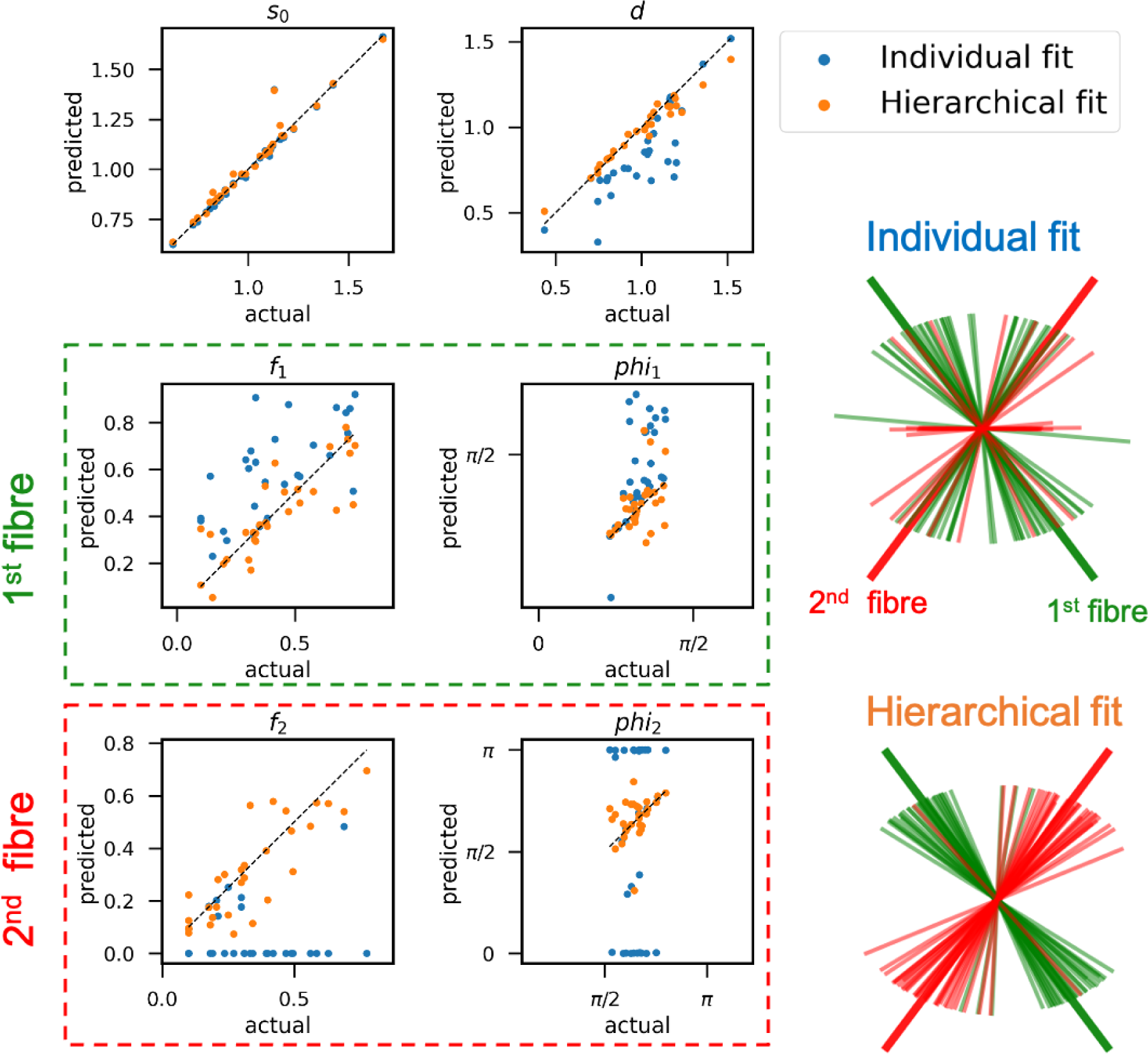
Scatter plots of estimated vs. actual parameters for the 2-fibre ball and stick model in a single voxel for 30 subjects using hierarchical and individual fits. For easier visualisation of the orientation parameters, we fixed the values of their spherical polar angle for all subjects to be 90 degrees and only showed estimates for the azimuthal angle phi. Each dot represents a subject, and the closer it is to the identity line (represented by the dashed line), the better the fit. The fibres in the individual fits are sorted by strength within each subject, while the fibres in the hierarchical fits are labelled according to the strength of the group-average fibre. For some subjects, the individual fit has swapped the values of f1 and f2 as well as the associated angles. In the lower-right corner, the estimated orientations of the sticks for individual fit and hierarchical fit are shown. The bold, elongated lines display the actual group average orientation. Despite the fact that the strength may have been in reverse order for some individuals, the hierarchical fit was able to label the sticks matching the correct labelling. In contrast, we observe a large amount of group variance in the individual fit method.

The results indicate that the hierarchical fit is more accurate for estimating the model parameters. In particular, the individual fitting approach failed to detect the weaker secondary fibre population in the majority of subjects (most of the second fibre signal fractions are zeros). The line plots show the estimated orientation of the sticks for each subject in both approaches. The individual fit has no clear clustered pattern whereas the hierarchical fit was able to assign the same label to corresponding sticks.

### White matter fixel template

The white matter fixel template contains the mean and variance of the ball-and-sticks model parameters for each white matter voxel, including the diffusivity, direction, and signal fraction of the fibres in that voxel. This provides a basis for comparing the signal fractions in different fibres over a broader sample of participants by fitting the ball-and-sticks model using the template as prior.

Figure 6 displays two sections of the template’s fixel orientations and two magnified areas of crossing fibre regions for better visual inspection. These orientations are calculated using the group average across subjects. Although there are no between-voxel spatial constraints, the orientations of the fixels exhibit good spatial continuity and conform to well-established anatomical features.

**Figure 6.**
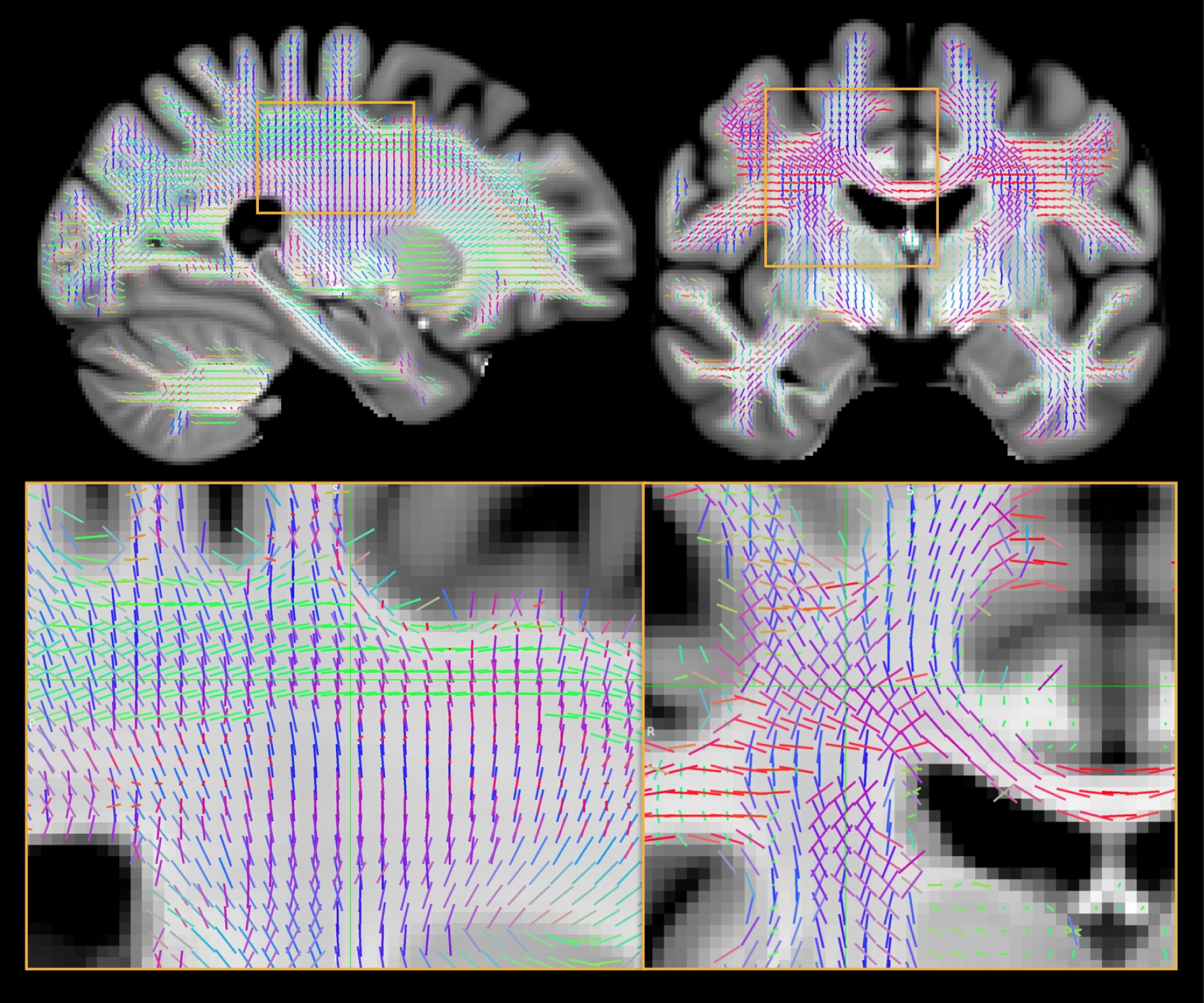
A sagittal and coronal slice of the white matter fixel template created using HCP diffusion data overlaid on the OMM1 T1-weighted image. The fixels that are present in each voxel are depicted by lines that are colour-coded according to their orientation (L-R: red, I-S: blue, A-P: green). The number of fixels is decided by thresholding the likelihood function improvement that is achieved by adding more sticks. Two sample patches of crossing fibre areas are magnified for better assessment. Although the technique does not involve regularisation across voxels, the fixels exhibit a high degree of spatial continuity. At the boundaries, we observe some fixels entering the grey matter radially.

Figure 7 compares the estimated fibre orientations for a sample white matter crossing fibre voxel with and without the hierarchical model. As a result of the subject-specific nature of the individual fit, existing fibre populations may not be oriented consistently across subjects, and some individuals may have a different number of fixels. Also, despite the fact that there is a high degree of consistency in the orientation of the fibre population between subjects, in some of them the relative strength of the fibre populations are swapped.

**Figure 7.**
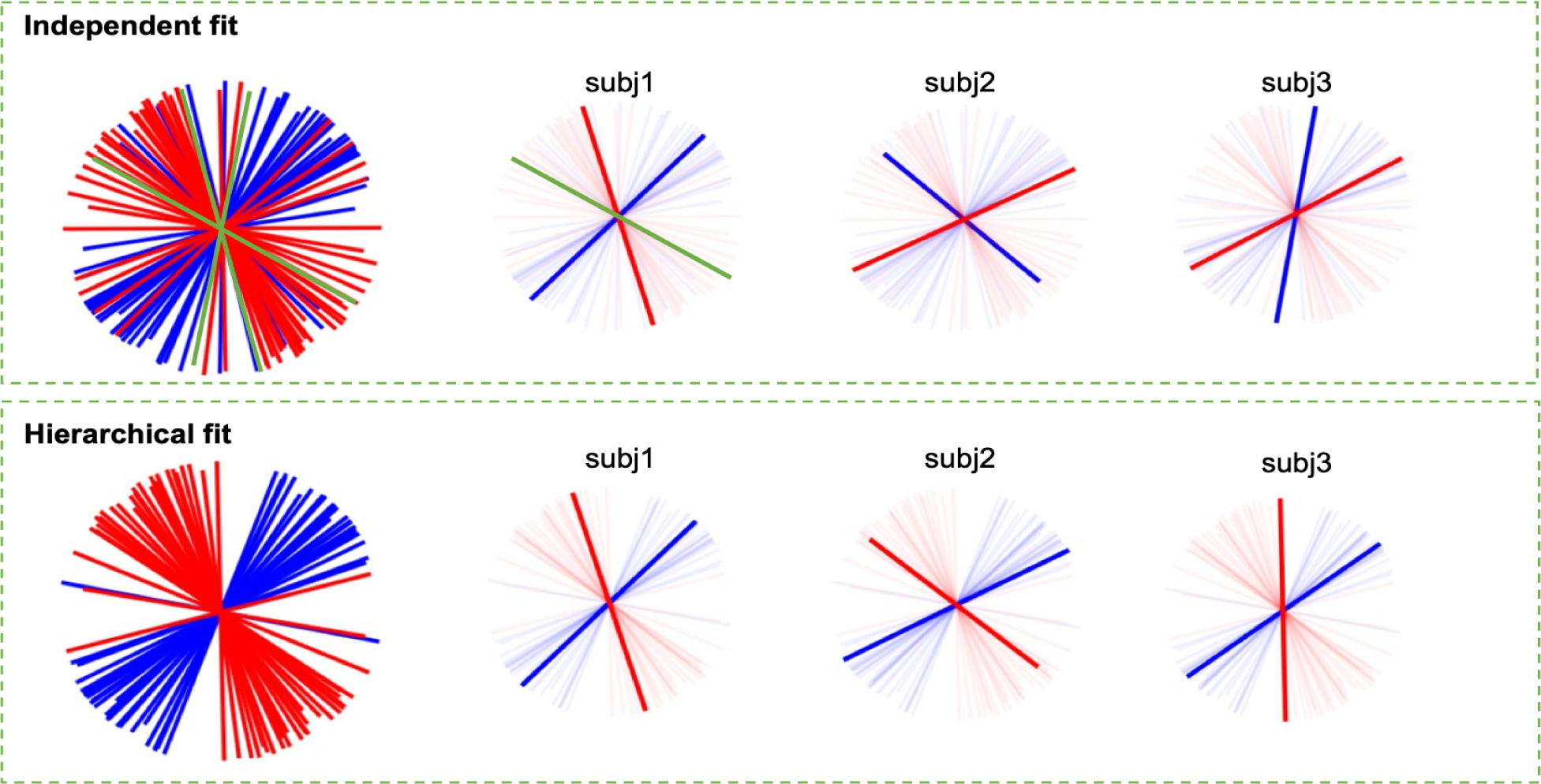
Estimated fixels in a sample voxel with crossing fibres for 30 subjects. The top panel shows aggregation of all subject fixels, along with three representative subjects. The first, second, and third fibres are represented by the colours blue, red, and green, respectively, in order of signal fraction. Various types of variability are observed across subjects: In this voxel in Subject 1, there are three distinct populations of fibres, in Subject 2, the first and second fixels are reversed compared to the other subjects, and in Subject 3, the fibres are oriented at an acute angle to one another. Any post hoc analysis to match fixels between subjects becomes challenging as a result of these differences. The hierarchical fit for the same data is shown in the lower panel. The labelling is based on the strength of the group average. In the aggregate plot, we can see that the fibres have organised into two distinct clusters. Similarly, the sample subjects demonstrate a strong correspondence with the template and between them.

The same fit is displayed in the lower panel, but this time the hierarchical model with 2 fibre populations was used. The results demonstrate that the fibres are aligned in two distinct groups and are matched between subjects. Furthermore, note that just the labelling has changed, and that the estimated orientations for each subject are extremely close to those from the individual fits. In other words, the hierarchical fit does not bias the orientations towards the group average and mostly only re-labels them.

Maps and a histogram of the cross-subject variation in orientation of the first fibre population (all white matter voxels) are shown in Figure 8 for both individual and hierarchical fits. The across subjects variability of the major fibre orientation in the individual fit is substantially higher than that for the hierarchical fit. This is due, in part, to the fact that certain subjects first and second (or third) fibre populations have switched places, and also because the hierarchical fit tends to pull orientations closer to the group average when data is not strongly supportive of a particular orientation (e.g., in outlier or noisy subjects). The maps depict a slice of the cross-subject dispersion estimated in the white matter voxels. In the hierarchical fit, any high variance resides close to the grey matter boundaries, where there is no strong and consistent fibre orientation structure.

**Figure 8.**
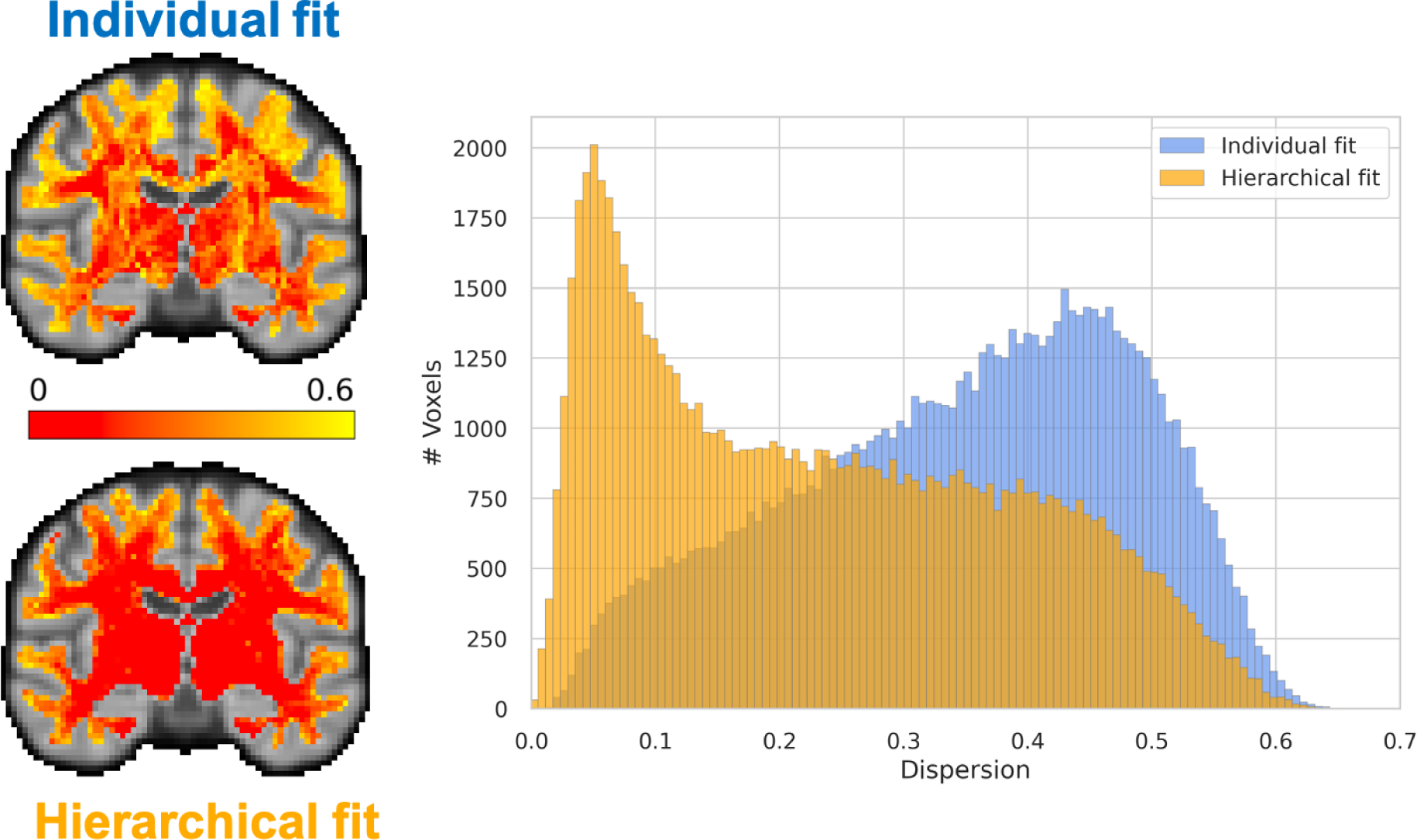
Variability of the primary fibre population across subjects with and without hierarchical fit: The histogram shows the distribution of across subject dispersion in all white matter voxels for both individual fits and hierarchical fit. The dispersion along the x-axis is quantified using the orientation dispersion index, as defined in the NODDI paper (H. Zhang, Schneider, et al. 2012). This index is computed by fitting a Watson distribution to the subject orientations, estimating the concentration parameter, and subsequently converting it to the orientation dispersion index. A noticeable reduction in cross-subject variability is observed, indicating a higher degree of alignment among the primary fibre population across subjects for the majority of voxels. This reduction can be attributed to both the hierarchical framework drawing subject parameters towards the group average and also better relabelling of fibres. The accompanying maps display dispersions for a single slice of the standard brain. In the hierarchical fit, it is evident that the majority of high dispersion voxels are located at the grey matter boundaries, while fibres in the deep white matter exhibit increased alignment across subjects. In contrast, this pattern is not observed in the individual fitting approach, especially at the crossing fibre regions.

The number of fibre populations in each white matter voxel is depicted in Figure 9 in example slices. The improvement in the likelihood function by adding an extra fibre population is the criterion for selecting the number of fibre populations in each voxel. The left histogram depicts the distribution of the difference in log-likelihood between models with two and one fibre population, summed across all participants. We apply a threshold to this histogram in order to identify voxels containing a single fibre population. The threshold is chosen by visually inspecting the maps to ensure a certain degree of symmetry across hemispheres and reasonable spatial homogeneity. The histograms on the top right displays the distribution of the log-likelihood difference between the 2 and 3 fibre models, but only for the voxels that are over the 1 fibre threshold. We apply a threshold to this histogram in order to identify voxels with two or three fibre populations.

**Figure 9.**
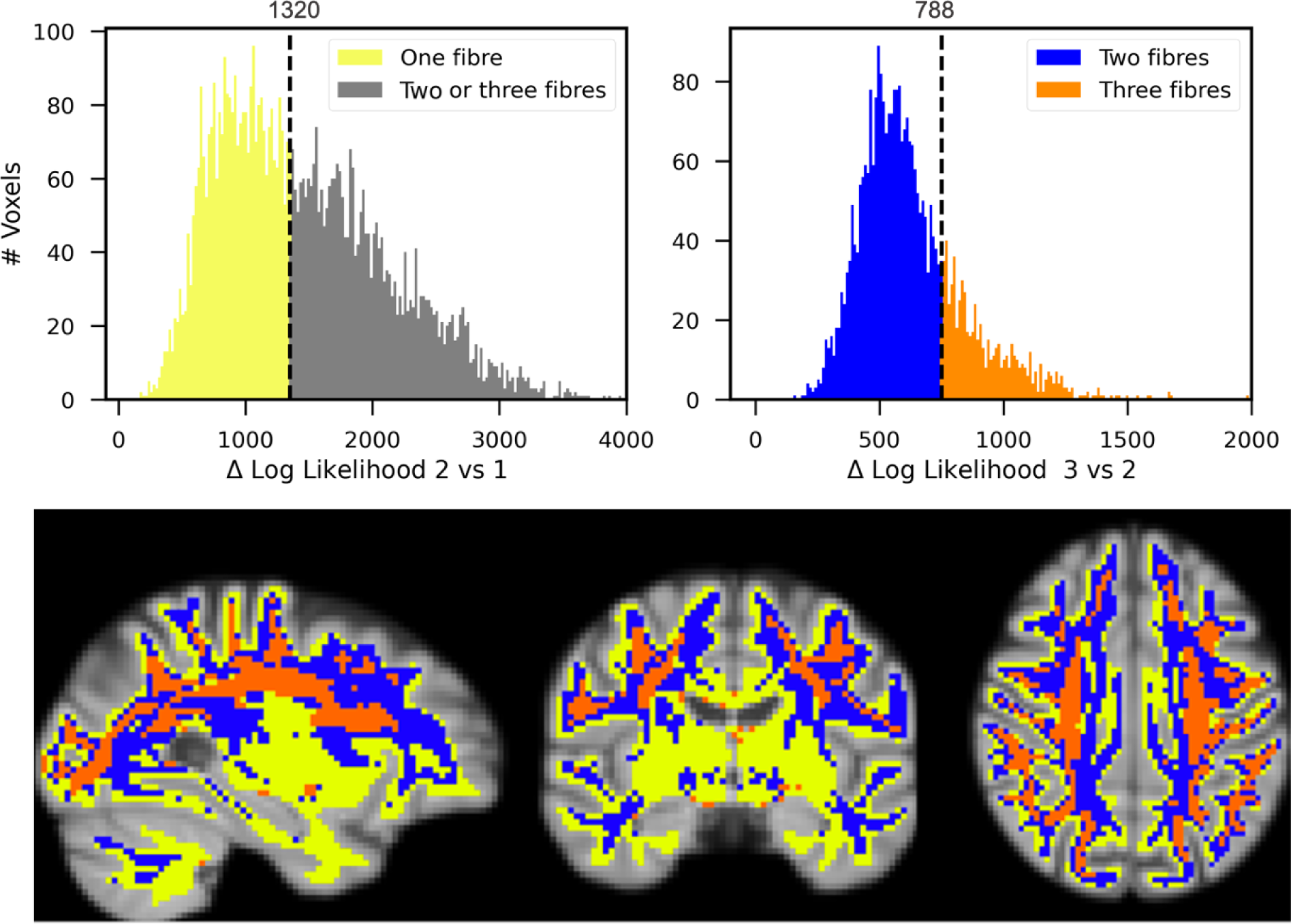
Determining the number of fixels present in each voxel. The histogram on the left displays the distribution (across the voxels in the white matter mask) of the increase in subject-averaged log likelihood from one to two fibres. We expect a better fit to data with more free parameters; hence all numbers are positive. To determine which voxels, contain only one fibre we threshold these differences. The threshold is chosen by visually inspecting the maps and taking into account across hemispheres symmetry and spatial homogeneity. The right histogram illustrates the improvement between two to three fibres exclusively for voxels that pass the criteria for one versus two models. This histogram is thresholded to indicate which voxels have two fixels and which have three fixels. The criteria for threshold is once more spatial homogeneity and cross-hemisphere symmetry. Maps depict a slicing through the template, with each voxel coloured according to the number of fixels it contains (yellow for 1, blue for 2, and orange for 3).

### Fixel based GLM

To illustrate how the fixel-based framework can be put to use for statistical analysis, we conducted a GLM analysis to assess the changes in white matter fibre tracts through ageing. For each subject, the HCP white matter fixel template was used as a prior to estimate fixel strength and orientation within each voxel.

Figure 10 depicts a section of white matter with fixels coloured according to the t-statistics associated with the age regressor from the GLM; With red denoting an increase in fibre signal fraction and blue denoting a reduction. To facilitate a better visual evaluation, a patch of crossing fibre areas is magnified. We observe both positive and negative changes in the region, with the majority of horizontal (anterior-posterior) fibres displaying positive changes and the majority of vertical (superior-inferior) fibres displaying negative changes. The histogram depicts the distribution of t-stats separated for horizontal and vertical fibres inside the patch. The t-stats for horizontal fibres are centred around zero, whereas the vertical population is skewed to the left.

**Figure 10.**
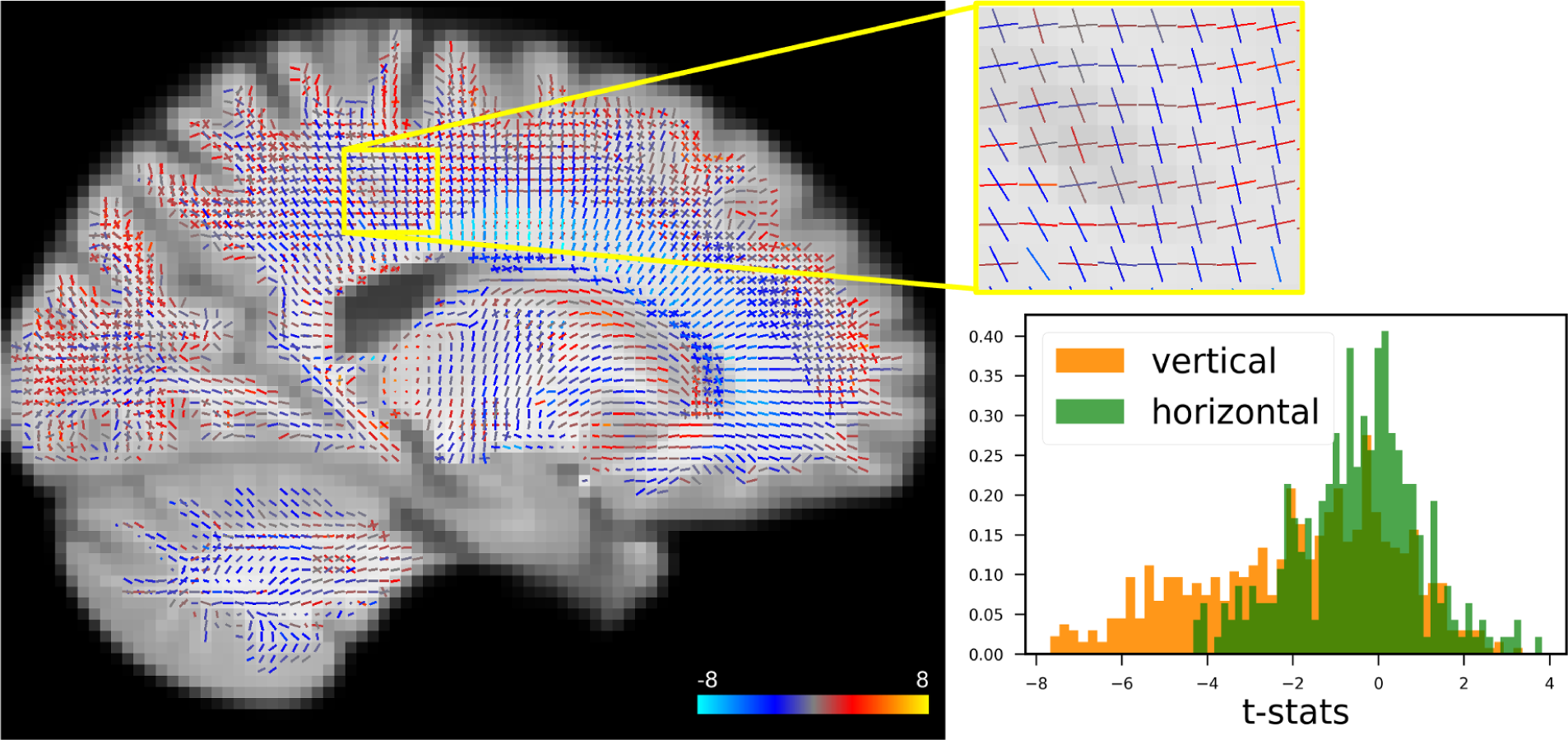
(Left) A sample slice of t-statistics for fixels. Colour of lines indicate correlation with age (blue for negative, red for positive correlations). (Right). Histogram of the t-statistics for horizontal and vertical fixels in the magnified region and 5 slices parallel to it. In this illustration, the distribution for vertical fibres is shown to be highly biased towards negative values, whereas the horizontal fibre distribution is centred around zero. In other words, on average, vertical (projection and commissural) tracts in this region degrade with age, whilst horizontal (association) tracts do not alter consistently.

Figure 11 displays the changes in signal fractions and fractional anisotropy for a representative deep white matter voxel as a function of age. The FA scatter plot trend indicates a significant rise in FA, which is sometimes interpreted as an increase in fibre integrity, when we would usually expect a decrease with ageing. Examining the fixel signal fractions helps us interpret this counter-intuitive result. The signal fractions for the three fibre populations in this voxel as a function of age are displayed in the scatter plot on the right. Rather than an increase in overall strength, the population of fibres with the lowest signal fraction is decreasing at a faster rate compared with the other two. This can explain the apparent increase in FA since anisotropy should increase, particularly when there is a large angle between this fibre population and the other two (see Figure 1).

**Figure 11.**
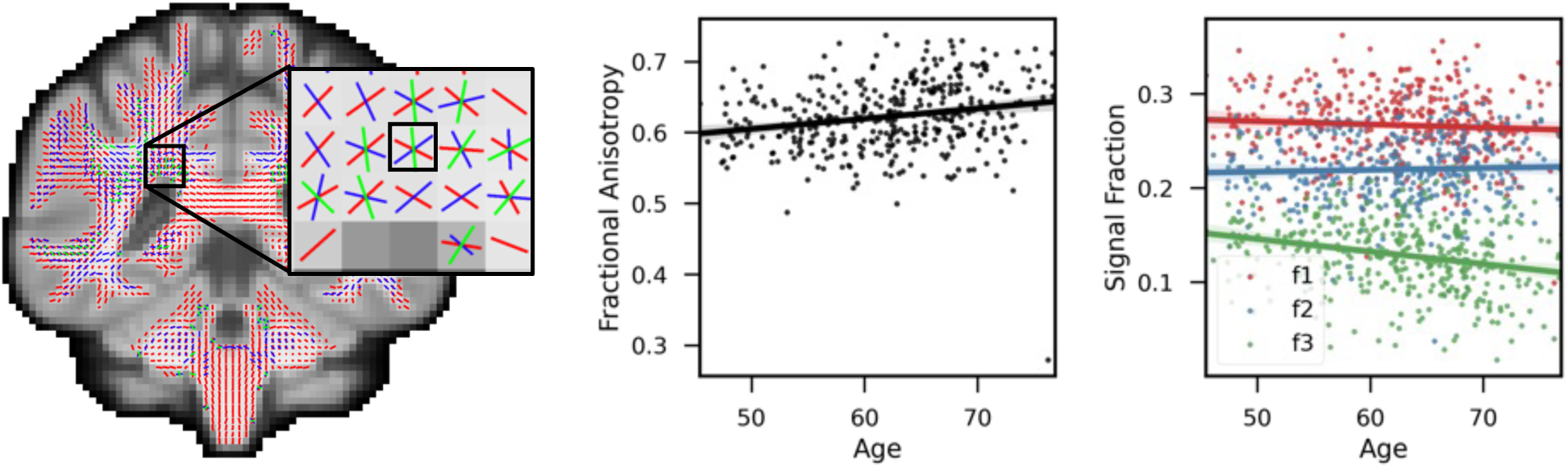
Comparison of FA and signal fractions in a sample voxel. Fractional anisotropy versus age for 400 subjects is shown in the scatter plot on the left (each dot represents a subject). In this voxel, the trend line (thick red line) suggests a positive FA-age correlation, which is conventionally attributed to an increase in fibre strength. The scatter plot on the right shows the signal proportions with trend lines for the three fibre populations present in this voxel (Red for first, Blue for second, and green for third fibre). While the first two fibre populations in this voxel changed very little with age, the third population decreased substantially with age. As a result, anisotropy increases because diffusion decreases along the direction of the changed fibre population relative to the other two.

Figure 12 depicts a map of fixels with significant changes (p<0.05) following multiple comparison correction with fixel-based threshold free cluster enhancement. Only negative changes in the vicinity of the corpus callosum survived the significance test.

**Figure 12.**
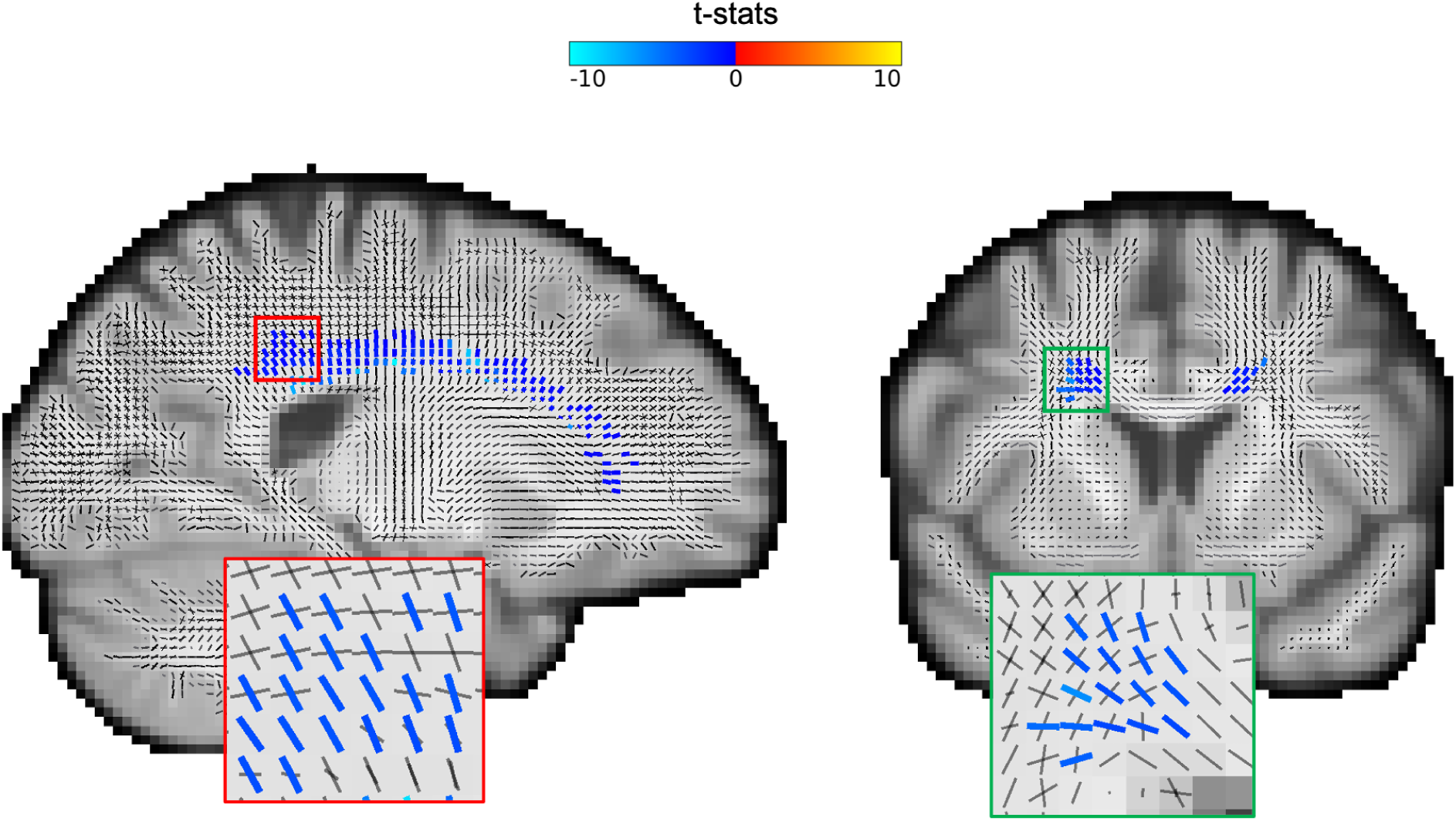
Statistically significant changes. Fixel-specific threshold-free cluster enhancement is applied to the t-stats derived from the GLM analysis on fixel signal fractions versus age (E = 1, H = 3). Permutation tests with 5000 repetitions on the TFCE values are used to estimate p-values for each fixel. The maps depict the significant fixels (colour-coded based on the t-statistics) with corrected p< 0.05. Only negative changes in signal fractions pass the statistical significance test, and they are all situated in the same tract (body of corpus callosum) in both hemispheres.

In Figure 13, we examine the changes with age in pairs of fixels forming crossing fibres in all voxels of the white matter. The figure displays the t-statistics of the first (x-axis) and second (y-axis) population of crossing fibres. Changes that occur in both populations of fibres and in the same direction (around the identity line in the scatter plots) represent coherent fixel changes, i.e. voxel-wise change, as opposed to changes that occur in other directions (away from the identity line) that are fixel specific changes. Significantly positive coherent changes (t-statistics in both fibres being more than 3) are shown by green dots in the scatter plot and displayed in the same colour on the maps. Voxels that exhibit substantial fixel specific changes (t-statistics difference is greater than 3 in either of the fibres) are identified by red colour and displayed as such on the maps. We find that most coherent positive changes occur along the grey matter boundaries, suggesting changes in grey/white partial volume with age, whereas most fixel specific changes occur in the deep white matter and likely capture fibre-specific changes in white matter organisation.

**Figure 13.**
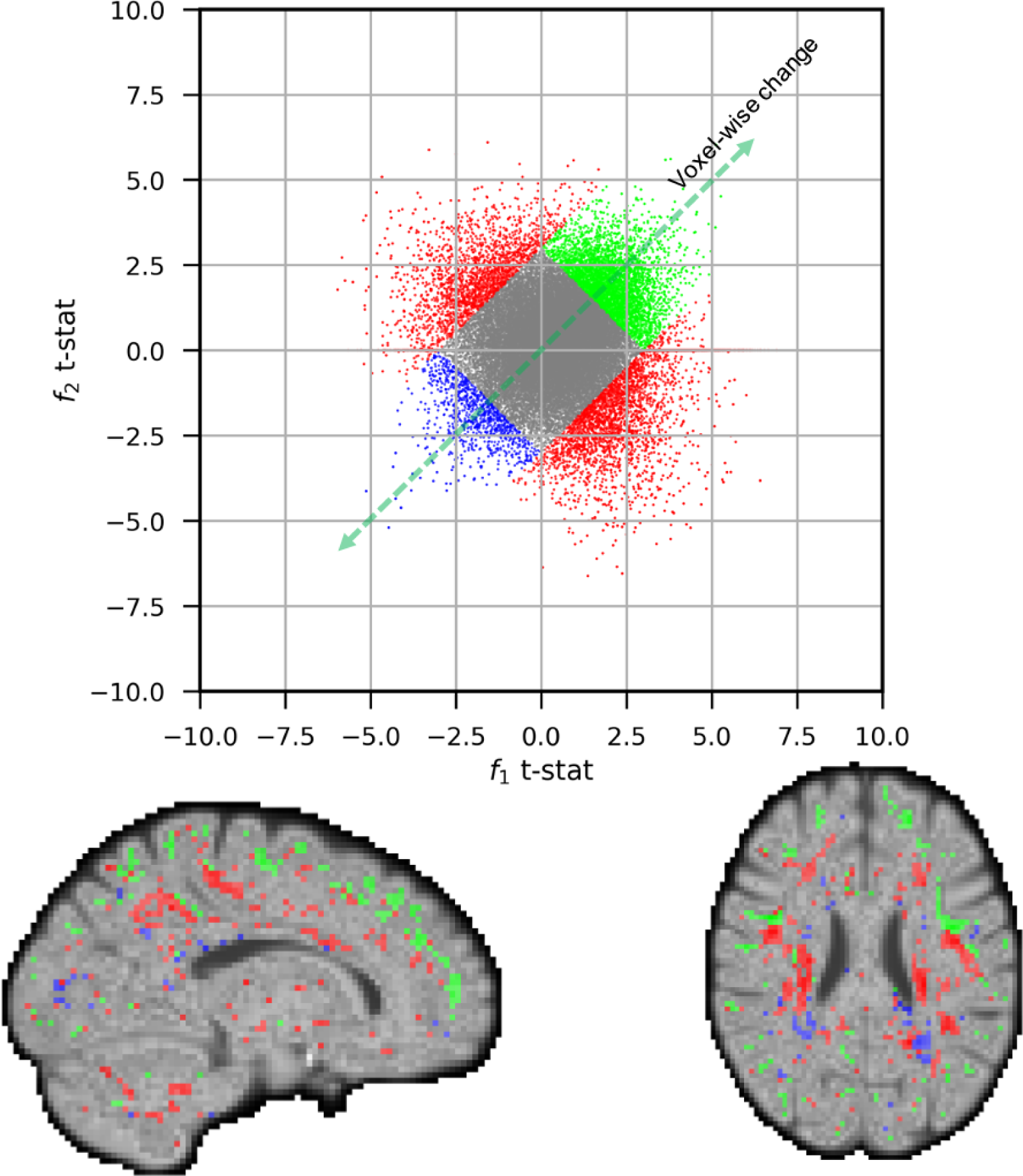
The scatter plot illustrates t-statistics for the the relationship between signal fraction and age for the first and second fibre populations across all white matter voxels. Each data point corresponds to a single voxel and axis represent the first and second fixel’s t-statistics. In certain cases, both fixels within a voxel exhibit the same changes, resulting in points clustered near the identity line (voxel-wise changes). Significant points of these instances (sum of t-values in both fixels being greater than 3 or lower than −3) are denoted in blue for negative changes and green for positive changes. Conversely, there are voxels where the fixel strengths change differently with age, shown by data points deviating from the identity line. Significant points with these changes (difference in t-values being greater than 3) are highlighted in red. The accompanying maps reflect these results using the same colour scheme as the scatter plot. Notably, the majority of coherent positive changes at the voxel level are concentrated near the grey matter boundaries, while specific fixel changes or coherent negative changes are primarily observed in deep white matter regions.

### SOFTWARE IMPLEMENTATION

WHIM(White matter HIerarchical Modelling) is an open-source software package and freely accessible from https://git.fmrib.ox.ac.uk/hossein/whim. WHIM is versatile and allows for the execution of full hierarchical inference, the generation of new templates, and the analysis of new data using available templates. WHIM will be integrated into new versions of FSL.

## DISCUSSION

In this study, we introduce a new approach for extracting fibre-specific measurements (fixels based metrics) from diffusion MRI data. We used a hierarchical framework to fit the ball-and-sticks model to data from multiple subjects that offers some advantages over an individual fitting approach. First, the cross-subject fibre labelling issue is inherently resolved by the hierarchical structure of the model. This eliminates the need for post hoc fibre matching, allowing us to directly compare fibre-specific parameters across individuals. The hierarchical structure also helps with robust parameter estimation for noisy subjects as it makes use of pooled data from multiple subjects. Natural expansions are viable in light of the hierarchical structure, such as hierarchically modelling subpopulations of individuals or employing a high-level model (such as age variables) as part of the inference process.

The hierarchical structure has some disadvantages as well. First and foremost, it is computationally more expensive than fitting the model to individuals separately. We addressed this issue by employing an EM algorithm, which converges within a few iterations and allows for both memory efficiency and convenient parallelisation across subjects. However, we note that once a template has been created, it can be utilised as a prior for fitting new subjects and would have the same computational costs as the individual fitting approach. Individual variations in the number of fixels within a voxel constitute another issue. Our method assumes that all subjects have the same number of fixels and is inflexible in this regard, but at least the hierarchical model enforces comparison of fixels with similar characteristics. Moreover, the hierarchical model requires accurate registration of white matter both in terms of location and local fibre orientation. Moreover, it is essential to note that the fixel template is designed for application to relatively normal brain tissue. That is to say, the template’s use on severely damaged tissue, where fibres have been significantly displaced or destroyed due to a tumour or injury, may produce unreliable results.

While the proposed framework is one option to extract fibre specific measurements, it is not the only one; the MRTRIX software (J-Donald Tournier et al. 2019) also has a very solid pipeline implemented that has been used in many studies. There are two key differences compared to our approach. Firstly, we estimate individual fibre orientations using the ball and sticks model while the other approach uses spherical deconvolution. Secondly, they solve the fibre assignment across subjects post hoc, after model fitting, whereas we perform it as part of the modelling process. Both methods provide proxy measures for fibre strength.

As of now, the diffusion tensor metrics such as FA and MD are the most frequently used measurements to examine the integrity of brain tissue. A GLM analysis for FA vs age reveals that a number of brain areas exhibit significant positive correlation with age. While a decrease of FA is often interpreted as reduced tract integrity, increases in FA are more difficult to interpret. In Figure 11, we provided a concrete example of real data demonstrating that a drop in the signal fraction of a crossing fibre in an example voxel in deep white matter is contributing to a rise in FA. Also, after applying the cluster corrections, the results of the fixel based GLM analysis show that all significant changes in the fibre strengths are negative. This would imply that the FA increases are confounded by the geometrical structure of the tracts and their relative alterations within each voxel.

### Fixel template creation and usage

We have applied the proposed methods on a few subjects from the HCP young adult dataset to produce a white matter fixel template. This template contains the probability distribution for the orientation of fibres present in each white matter voxel in the Oxford Multimodal Template 1 (OMM-1), which can be used in the future for fibre-specific statistical analysis. It may be essential to create a new template if a cohort-specific template is required (e.g., for disease or a different age range), if using a different base diffusion model (other than ball-and-sticks), or if using a structural template other than OMM-1.

It is worth re-emphasising that whilst the hierarchical model can be fitted to a cohort of subjects using our EM algorithm, the resulting group parameters constitute a template that can be employed in its own right as a prior to fit additional subjects that are not part of the same cohort. This is not only useful for computational reasons. It can also benefit in fitting lower quality data than those used to create the template. For instance, it is well established that in diffusion modelling, there is a tension between acquiring multiple shells to have a better estimate of the diffusion parameters (e.g., ADC and kurtosis etc.) and acquiring multiple directions to have more accurate crossing fibre modelling. Most datasets compromise on one or the other. The template can be created using high-quality, multi-shell and multi-directions data, such as the HCP data, and then used as a prior to constrain the analysis of lower quality data. We have used exactly this approach here, with a template created using HCP, and deployed to analyse the UK Biobank data. Nevertheless, when adapting the template to datasets obtained through different protocols, it’s advisable to exclude the priors on diffusivities. This is because these measurements might be influenced by the specifics of the acquisition protocols and the strength of the gradients used.

In this study, the hierarchical structure was applied to all parameters of the ball and sticks model, excluding the baseline signal (S0) parameter. This decision was made due to the potential for artefacts, such as bias field, to induce intrasubject variability in S0 with a complicated cross-subject distribution. Incorporating a parameter into the hierarchy can help to regularise its estimates especially in subjects with noisy data, but it can also induce biases. It is not required to include all parameters in the hierarchy; for example, signal fractions might be left out to avoid potential biases. However, stick orientations must always be maintained in the hierarchy to enable consistent labelling between subjects.

In order to determine how many fixels are present in each voxel, we have relied on the increase in log-likelihood as a measure of goodness of fit. In determining an appropriate cut-off for this metric, we take into account the spatial coherence in the produced maps and whether they are consistent with the information we currently have on the total number of tracts in each region. To determine the thresholds, one may also use metrics such as AIC and BIC. However, there is always an arbitrary trade-off between the number of free parameters in the model and its goodness of fit. In any case, the thresholds are fairly flexible and can be re-adjusted with minimal effort if there is a compelling reason to do so. Additionally, automated methods for determining the number of fibers may be developed and readily integrated into this framework in the future.

In addition to the free parameters that are fitted using data, the hierarchical model also includes user-set hyperparameters (α_g_, α*_n_*) that constrain the prior on the group variance and the noise variance. Having a high α_g_ imposes a greater penalty on group variances to approach zero, hence increasing the chance to favour individual differences. A high α*_n_* increases the penalty for estimating high noise variance. Changing either of these parameters within a suitable range, however, should not significantly alter the model’s behaviour.

### Future Directions

The hierarchical structure can be enhanced further in a number of ways. The template can be created in a nested hierarchical structure for multiple groups of subjects, and comparisons can then be made between group parameters. Another modification is to employ a regression model to incorporate independent variables (such as age) into the hierarchical structure. Moreover, in order to better handle noisy subjects, one can additionally include the uncertainty in subject parameter estimates into the model and convert it to a mixed-effect analysis.

The fixel template is created by employing the ball and sticks model to extract the orientations of the fibre tracts within each voxel. The ball and stick model were chosen as it is a well-tested model designed to robustly extract crossing fibres in the brains white matter. However, this does not imply that the tract specific metrics are restricted to these model parameters. The presented template can be used as a starting point for developing more complex and specific biophysical models to fit the data. For instance, the template can be used to fit fibre dispersion or multi-compartment models (e.g., NODDI (H. Zhang, Schneider, et al. 2012)) with crossing fibres by utilising the information on the number and orientation of fibres in each voxel. The parameters of those biophysical models can subsequently be employed as fixel-specific measurements that are compared across subjects.

As an illustration of the proposed method, we have employed a GLM framework to conduct fixel-specific statistical analysis in this study. The steps for performing such an analysis and the results to anticipate are laid out in detail here. Age has been used as an independent variable to clarify usage, but we hope that the tool will be useful to explore fibre-specific alterations as a function of disease or other biological processes. Moreover, it is possible to explore fiber-specific changes in the brain by integrating these measurements with data-driven techniques like Independent Component Analysis (ICA) or Canonical Correlation Analysis (CCA). To achieve this, one can create a fixel-by-subject matrix by concatenating all the fixels’ data, and then apply matrix factorization algorithms to estimate patterns of variation within the fixels. This approach helps identify modes of variation in the population, providing insights into how fixel strength varies across subjects. Additionally, the loadings obtained from this analysis can be valuable for identifying potentially interesting and meaningful individual differences. Moreover, tractography approaches can be applied to the template to reconstruct the major white matter tracts, enabling the across subjects comparison of brain structure along the tracts.

In order to increase the power of statistical tests for fixel-based data, here we present a modified version of threshold free cluster enhancement. Similar to the original TFCE technique, this method utilises two hyperparameters (E and H) to adjust how heavily cluster extent and height play towards promoting significant clusters. In this study we have used the original method’s (E=3, H=2) values for these parameters, which have been shown through simulations to be appropriate for voxel-wise data. Future development may involve adjusting them to accommodate for the different neighbourhood structure of fixel-based data compared to that of voxel-based data.

